# Minimum effort simulations of split-belt treadmill walking exploit asymmetry to reduce metabolic energy expenditure

**DOI:** 10.1101/2022.08.15.503979

**Authors:** Mark Price, Meghan E. Huber, Wouter Hoogkamer

## Abstract

Walking on a split-belt treadmill elicits an adaptation response that changes the baseline step length asymmetry of the walker. The underlying causes of this adaptation, however, are difficult to determine. It has been proposed that effort minimization may drive this adaptation, based on the idea that adopting longer steps on the fast belt, or positive step length asymmetry (SLA), can cause the treadmill to exert net-positive mechanical work on a bipedal walker. However, humans walking on split-belt treadmills have not been observed to reproduce this behavior when allowed to freely adapt. To determine if an energy minimization motor control strategy would result in experimentally observed adaptation patterns, we conducted simulations of walking on different combinations of belt speeds with a human musculoskeletal model which minimized muscle effort. The model adopted increasing amounts of positive step length asymmetry and decreased its net metabolic rate with increasing belt speed asymmetry, up to +25.6% SLA and −14.3% metabolic rate at a 3:1 belt speed ratio, relative to tied-belt walking. These gains were primarily enabled by an increase of braking work and a reduction of propulsion work on the fast belt. The results suggest that a purely energy minimization driven split belt walking strategy would involve substantial positive SLA, and that the lack of this characteristic in human behavior points to additional factors influencing the motor control strategy, such as aversion to excessive joint loads, asymmetry, or instability.

**New & Noteworthy:** Behavioral observations of split-belt treadmill adaptation have been inconclusive toward its underlying causes. To estimate gait patterns when driven exclusively by one of these possible causes, we simulated split-belt walking with a musculoskeletal model which minimized its energy cost. Our model took significantly longer steps on the fast belt and reduced its metabolic rate below tied-belt walking, unlike experimental observations. This suggests that asymmetry is energetically optimal, but human adaptation involves additional factors.

## Introduction

Over the past decade, split-belt treadmill training has emerged as a promising form of physical therapy to reduce gait asymmetries in neurological patients as well as a paradigm to study locomotor adaptation. During split-belt treadmill walking, the belts under each leg run at different speeds. For healthy individuals, this initially induces an asymmetry in some spatiotemporal patterns between the two legs (most notably step length), but over time, symmetry in these patterns returns (1). When the belt speeds are subsequently set to the same speed, an aftereffect is observed – an opposite asymmetry emerges but quickly disappears (1). To rehabilitate individuals with asymmetric gait, the belt speeds are controlled to exaggerate the asymmetry in step length (2). Unlike in healthy individuals, the aftereffect upon returning the belts to the same speed results in more symmetric gait for these individuals (2). While this aftereffect is also short-lived and individuals revert to their asymmetric gait, repetitive bouts of this split-belt treadmill training can reportedly lead to longer-term improvements in gait symmetry (3).

Traditionally, the change in human gait behavior during split-belt treadmill has been attributed to neuromotor adaptation (1, 4). It is often hypothesized that the nervous system considers inter-leg asymmetry an error that it drives to zero (i.e., towards symmetry) (5). Adaptation caused by error-based learning has been evidenced in motor tasks aside from split-belt treadmill walking and is generally considered to be a prototypical motor learning mechanism (6).

It has also been hypothesized that split belt treadmill adaptation patterns are driven by minimization of the energy cost of locomotion (7–9). Split-belt treadmills theoretically can exert net positive work on a user walking at constant asymmetrical belt speeds if the user adopts a positive step length asymmetry (SLA); greater step lengths with heel strike more forward of the body center of mass on the fast belt than on the slow belt (7). When initially presented with belt speed asymmetry, humans typically walk with negative step length asymmetry (longer step lengths at heel-strike on the slow belt than on the fast belt), but over time, they typically adjust to nearly symmetrical step lengths (9). Furthermore, it has been demonstrated that humans, when provided with guided experience of positive step length asymmetry gait patterns, tend to self-select a gait pattern with positive step length asymmetry (7). It has also been observed that humans, when allowed to adapt for a sufficiently long time (i.e. 45 minutes), will naturally adapt to positive step length asymmetry (8). There is conflicting evidence for this claim, however (10, 11). For instance, experimental participants tend to self-select a step length asymmetry that is smaller than the energetic optimum (10) and are unable to use the split belt treadmill to reduce the metabolic energy cost of walking below that of normal walking at the averaged speed between the belts, even in cases where the treadmill performs net positive mechanical work on the human (7, 8, 10, 11).

Overall, it is difficult to distinguish between motor control objectives and decouple them from the natural system dynamics from behavioral data alone (12). Optimal control simulations provide a framework for associating movement patterns with specific motor control objectives by optimizing the behavior of a transparent model according to those objectives (13). Prior work has most commonly used this approach to solve for energetically optimal gait patterns, such that resulting biomechanics can be fully explained by energy minimization within established constraints. This approach has provided insight into optimality principles underlying human gait (14–16), reproduced gait deficiencies (17–19), and demonstrated the potential outcomes of device-assisted gait (20–23). For split-belt treadmill walking, optimal control simulations can be used to determine if effort minimization can explain behavioral adaptation patterns, and if human musculoskeletal models can exploit belt speed asymmetry to reduce the metabolic cost of walking by increasing step length asymmetry.

In this paper, we simulated split belt treadmill walking by minimizing muscle effort and compared the results to experimentally observed gait patterns after adaptation. We hypothesized that minimum effort simulations of gait would display positive step length asymmetry to utilize the belt speed asymmetry to reduce muscle effort.

## Methods

### Model Description

We created a 2D musculoskeletal gait model by modifying the “2Dgait” model in OpenSim 4.3 (24) to include individually controlled treadmill belts for each foot, as depicted in Figure 1. The model consisted of trunk, pelvis, thigh, shank, foot, and walking platform rigid bodies with a total of 12 degrees of freedom in the sagittal plane. Nine pairs of bilateral “DeGrooteFregly2016” Hill-type muscles (25) actuated the lower limb joints, and ideal coordinate actuators actuated the lumbar joint angle and the walking platform anterior-posterior position. The muscle models in this set of simulations ignored tendon compliance, effectively assuming that the tendons behave like rigid links. The human model mass was 62.0 kg, and the height was 1.64 m.

**Figure 1.**
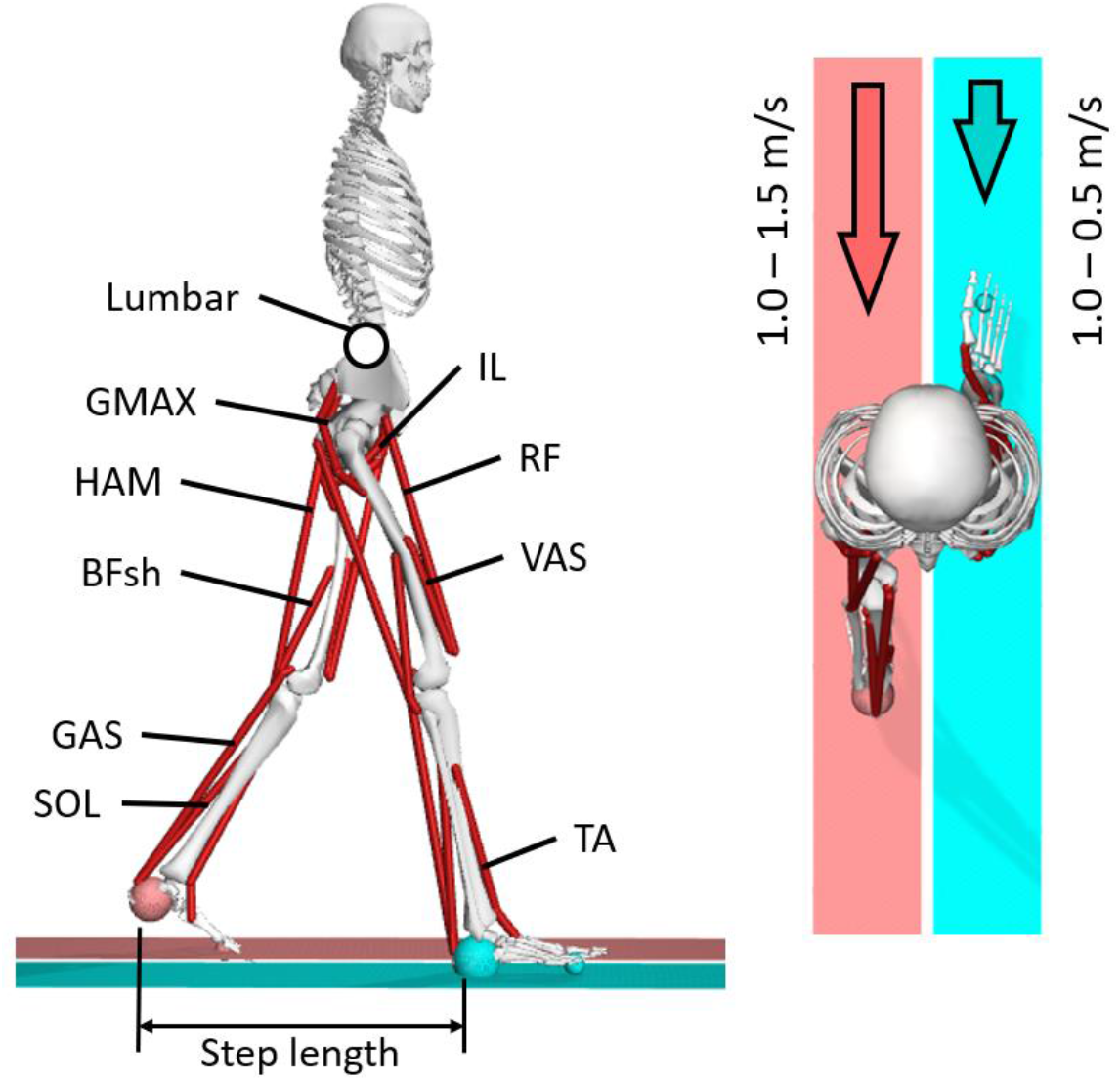
Two-dimensional musculoskeletal model for gait on a split-belt treadmill and asymmetric belt speed conditions. The legs are controlled by a set of 9 Hill-type muscle models on each side, for a total of 18 muscles. The lumbar joint is controlled by an ideal coordinate actuator which is not included in the energy cost minimization or the calculation of metabolic rate.

Ground contact was represented by spherical Hunt-Crossley contact elements located at the heel and ball of each foot and an infinite length contact plane located on the surface of each walking platform. Foot-ground contact pairs were established separately for left and right sides so that each belt only affected the corresponding foot. The platforms were constrained to a constant velocity relative to the global coordinate system to simulate the treadmill belts of a split-belt treadmill. The platforms each had a mass of 1000 kg to (1) reduce the active control adjustment required to maintain constant speed and (2) allow the average displacement of the treadmill belts to represent the change in the overall model center of mass position in the optimization cost function.

We generated simulations of periodic strides using a direct collocation optimal control method implemented through the Moco 1.1.0 package (26) included with OpenSim 4.3. Each stride was discretized into 100 mesh intervals and constrained to require realistic joint ranges of motion and identical start and end model states, with the exclusion of the belt positions. The objective function was described by Equation 1

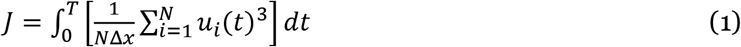

where *u_i_*, is the excitation of actuator *i* at time *t, N* is the total number of actuators in the model, and Δ*x* is the horizontal displacement of the model center of mass over one stride, which is in this case approximately the average backward displacement of the treadmill belts. This objective function approximates a muscle fatigue minimization approach in that large activation peaks in individual muscles are penalized more than with an aggregate metabolic cost of transport calculation, and competently reproduces level, steady-state walking overground (14)

#### Simulation conditions

We simulated periodic strides with six belt speed ratios ranging from 1:1 to 3:1, with the average belt speed remaining 1.0 m/s for each condition. We adjusted belt speeds in 0.1 m/s intervals from 1.0 to 1.5 m/s for the fast belt and 1.0 to 0.5 m/s for the slow belt.

### Analysis of Simulated Gait

#### Step length asymmetry

The gait cycle is comprised by two steps, one from heel-strike on the fast belt to heel-strike on the slow belt (*s→f*), and the other from heel-strike on the slow belt to heel-strike on the fast belt (*f→s*). We calculated step length as the distance between the left and right calcaneus coordinate frames at heel strike for each step, defined as the vertical ground reaction force rising above 25 N. *SL_s→f_* is defined as the step length at heel strike at the end of the *s→f* step, and *SL_f→s_* is defined as the step length at heel strike at the end of the *f→s* step. As common in split-belt treadmill walking studies (7, 9, 27), we calculated the step length asymmetry ratio as the difference between step lengths divided by the combined stride length:

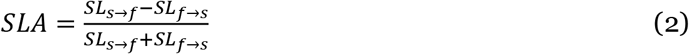

#### Step time asymmetry

We calculated step time where *t_s→f_* is defined as the duration of the *s→f* step from heel-strike to heel-strike, and *t_f→s_* is defined as the duration of the *f→s* step from heel-strike to heel-strike. The step time asymmetry ratio is defined by:

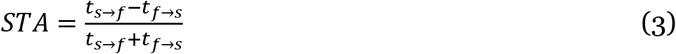

#### Foot placement

We defined the foot placement as the anterior-posterior position of the calcaneus coordinate frame with respect to the pelvis center of mass (approximately the human model center of mass). We calculated these measures for both leading and trailing limbs at heel-strike on each belt.

#### hGRFvs. limb angle

A proportional relationship between limb angle and horizontal ground reaction force (hGRF) magnitude is a fundamental assumption behind the simple conceptual model that produces positive net work transfer from a split belt treadmill to a human walking with positive step length asymmetry and mostly pendular mechanics. We compared peak horizontal ground reaction forces (hGRF) with the corresponding limb angle to determine if this assumption holds for a more complex model. We obtained the ground reaction forces from the foot-ground contact model interaction forces. We calculated the peak positive and negative horizontal ground reaction force (hGRF) for each leg and each belt speed condition to determine the effect of foot placement relative to the center of mass on horizontal ground reaction force magnitude. We defined the corresponding limb angle as the angle between the global vertical axis and a line drawn through the pelvis center of mass and the calcaneus limb segment. Although peak load should theoretically correspond temporally with peak limb angle, we also calculated the total braking and propulsive hGRF impulse magnitudes for each limb. Peak hGRF at the leading limb does not necessarily cancel with peak hGRF at the trailing limb, whereas propulsive and braking impulse must cancel across the gait cycle for the body not to drift forward or backward on the treadmill relative to a global reference frame.

#### Mechanical power

We calculated the propulsive, braking, and net power exerted by the legs as well as the power exerted by the treadmill belts on the body. As in (7), instantaneous power exerted by the legs is composed of power exerted on the body, defined as the dot product of the ground reaction force and the body center of mass velocity, and power exerted on the treadmill belts, defined as the dot product of the negative ground reaction force and the belt velocity. Positive and negative power components were calculated by separately integrating the positive and negative power regions with respect to time to obtain the positive and negative work components per stride, then dividing by the stride duration. Power exerted by the belts is defined as the dot product of the ground reaction force and the belt velocity for each belt.

#### Metabolic power

We calculated the metabolic rate for each muscle and in aggregate using the model proposed in (28), assuming a specific tension of 60 N/cm^2^ for all muscles and a basal metabolic rate of 1.0 W/kg.

All comparisons are made between the results obtained for the 1:1 and 3:1 belt speed ratios unless otherwise specified.

## Results

### Step length asymmetry

**Energy-optimal SLA increased with increasing belt speed asymmetry.** Step length asymmetry increased from 0 to +25.6% as the belt speed ratio increased (Figure 2a-b). The magnitude of step length asymmetry observed in the simulated model was larger than that experimentally found to be an energetic optimum (7) and larger than humans have previously selected (7, 8).

**Figure 2.**
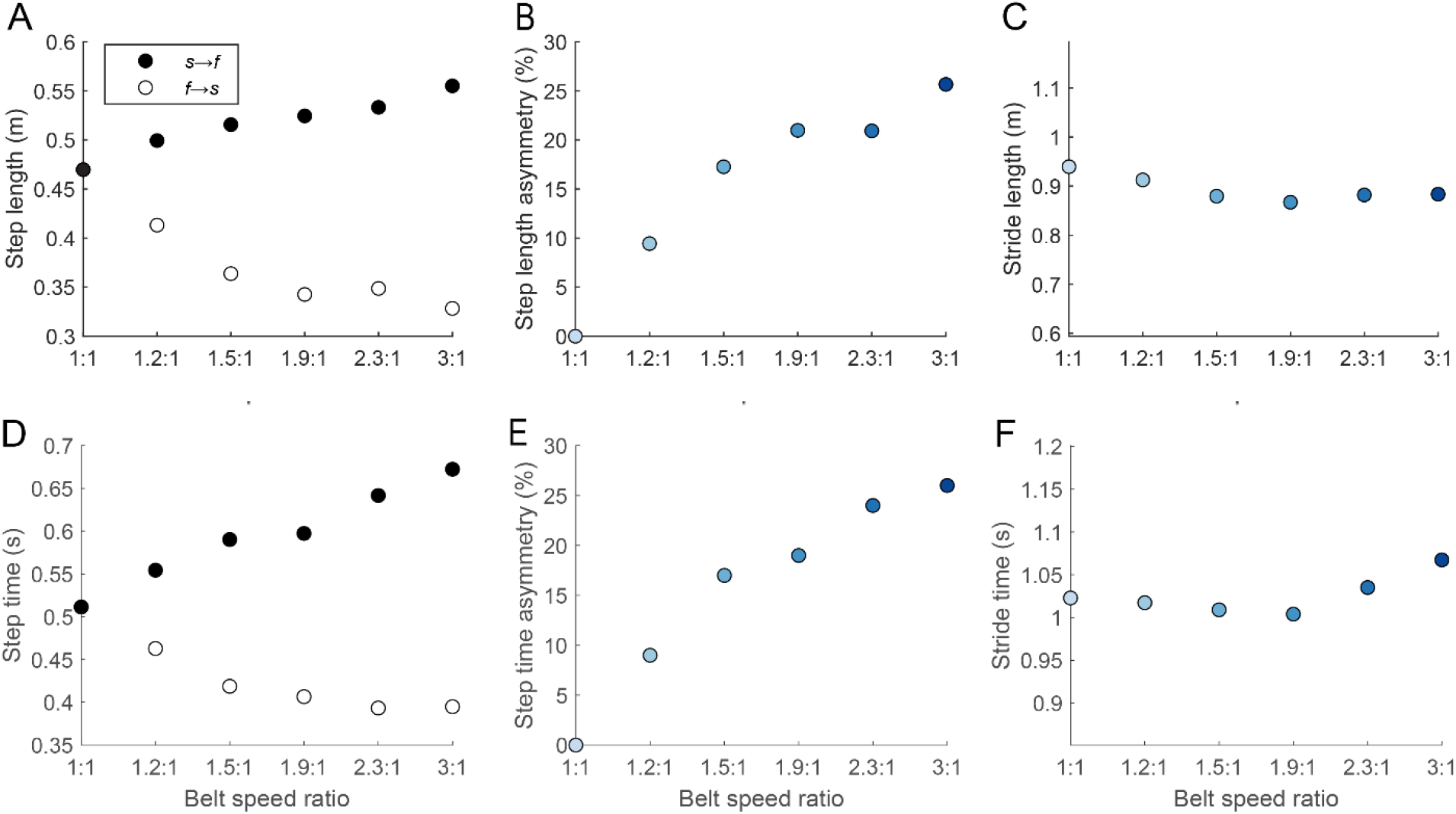
Top row: (A) Individual step lengths, (B) step length asymmetry, and (C) overall stride length vs. belt speed ratio. Bottom row: (D) Individual step times, (E) step time asymmetry, and (F) overall stride time vs. belt speed ratio.

### Step time asymmetry

**Energy-optimal STA decreased with increasing belt speed asymmetry.** Step time asymmetry increased from 0 to +26.0% as the belt speed ratio increased (Figure 2d-e). These changes are similar in magnitude to the changes observed in experimental studies (10).

### Total stride length and stride time

**Fluctuation in overall stride length and stride time was minor in comparison to the changes in symmetry.** Stride length decreased by 6.0% (Figure 2c) and stride time increased by 4.4% (Figure 2f) as the belt speed ratio increased. The minimum stride length occurred for a 1.9:1 belt speed ratio (belt speeds of 1.3 and 0.7 m/s), reaching a 7.7% decrease from the stride time at a 1:1 belt speed ratio. In recent experiments, stride length has been shown to decrease with increasing belt speed asymmetry though stride time showed little change (11).

### Foot placement

**Changes in step length asymmetry are driven by leading foot placement.** The change in step lengths can mostly be accounted for by the leading foot position for both sides (Figure 3). On the fast belt, the leading foot heel-strike position increased by +20.8% in front of the center of mass and the trailing foot position increased by 14.3% behind the center of mass on the opposite belt (Figure 3a). On the slow belt, the lead foot heel-strike position decreased by −54.5% in front of the center of mass while the trailing foot position on the opposite belt ultimately increased by 5.3% behind the center of mass, despite an initial decrease at lower belt speed ratios. Contrary to the rigid-leg simple model (Sanchez et al. 2019), step length asymmetry does not result in the trailing foot position shifting in the same direction as the leading foot position on the opposite belt (i.e., increasing or decreasing together in distance from the center of mass) for belt speed asymmetry ratios larger than 1.5:1.

**Figure 3.**
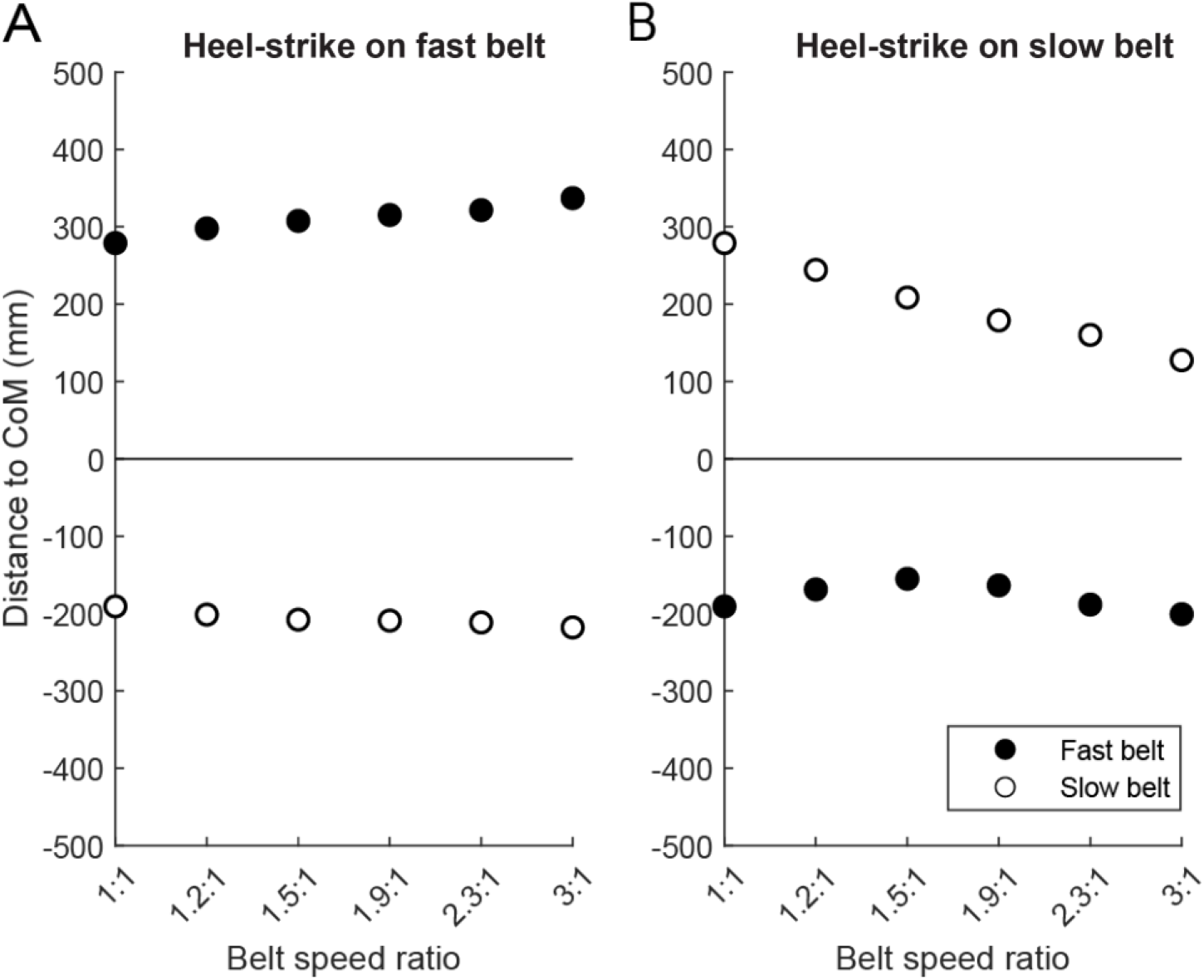
Heel distance from body center of mass for both leading and trailing legs at heel-strike (A) on the fast belt and (B) on the slow belt vs. belt speed ratio.

As has been observed elsewhere (10, 29), the constraints imposed by the treadmill create a coupling relationship between stride time, step length, step time, and foot placement for a given set of belt speeds. The simulation does not majorly alter stride time, but it does modulate the other three parameters to a much greater degree than has been observed in human experiments to reach a minimum metabolic effort solution.

### hGRFvs. limb angle

**Changes in horizontal ground reaction force depend on leading limb angle but do not vary proportionally with trailing limb angle.** Peak braking hGRF increased by 491 N on the fast belt at heel strike coupled with a modest increase in limb angle (+3.5°), while peak braking hGRF also increased at heel strike on the slow belt (46 N) despite a much larger opposing change in limb angle (−9.1°) (Figure 4a-b). Additionally, the change in peak push-off hGRF changes inversely with peak trailing limb angle in both limbs, counter to models which approximate the legs as pistons which make point contact with the ground, as used in (7). These results suggest that peak hGRF may be more dependent on belt speed than limb posture due to increasing belt speed necessitating a higher energy collision at heel-strike.

**Figure 4.**
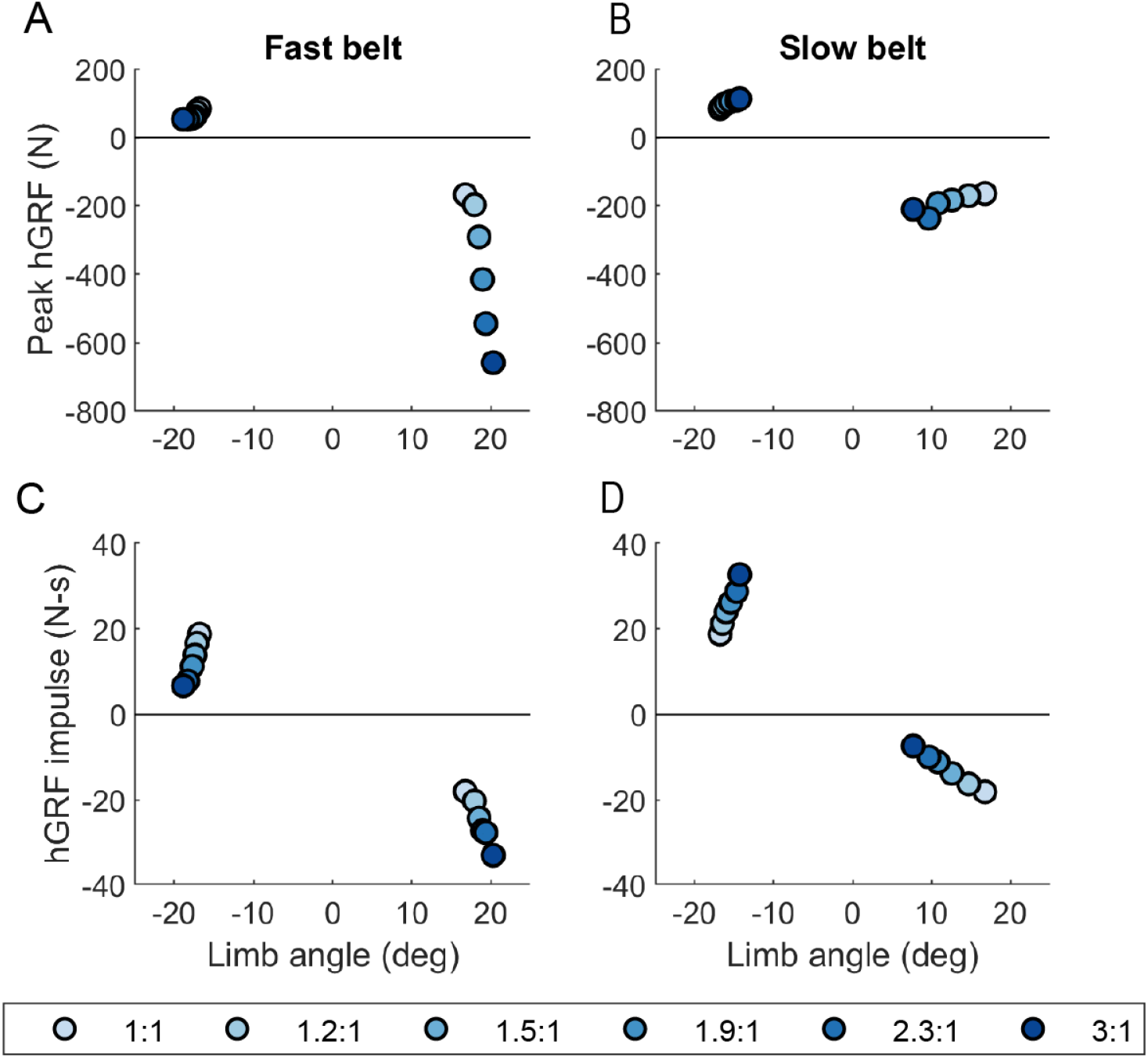
Peak horizontal ground reaction force during both heel-strike and push-off vs. the corresponding limb angle relative to vertical (A) on the fast belt and (B) on the slow belt. Total positive and negative contributions to horizontal impulse vs. limb angle at heel-strike and push-off (C) on the fast belt and (D) on the slow belt.

Propulsive and braking impulse changed inversely between limbs as the belt speed ratio changed (Figure 4c-d). The change in braking impulse corresponds proportionally with peak limb flexion angle for collision impulse for both legs, increasing by 15 N-s on the fast leg and decreasing by 11 N-s on the slow leg. This behavior is as expected by the simple dynamic model with pendular mechanics. However, as with the peak loads, propulsion impulse varied inversely with peak limb extension angle for both legs. Propulsion impulse decreased by 12 N-s on the fast belt and increased by 14 N-s on the slow belt as belt speed asymmetry increased, which agrees with the overall trend predicted by the simple dynamic model despite the lack of agreement on the relationship to horizontal load to limb angle. These results suggest that pendular mechanics apply for collision but not for propulsion, and that the model exploits this fact to shift most of the braking impulse to the fast belt and the propulsive impulse to the slow belt without requiring a substantial rearward foot placement on the slow belt.

### Mechanical power

**Asymmetrical belt speeds cause the treadmill to exert net positive work on the human for simulated gait.** The model further demonstrates that increasing belt speed asymmetry can be exploited to make the treadmill perform net positive work on the human per gait cycle. As observed in experiments and predicted by simple dynamical models, the human and treadmill perform approximately zero net work on each other at symmetrical belt speeds, but mechanical energy flows from the treadmill to the human at asymmetrical belt speeds, primarily through a sharp increase in mechanical power absorbed by the human model at heel-strike on the fast belt (Figure 5). Peak negative power performed by the fast leg increased in magnitude by 797 W as belt speed asymmetry increased. Peak negative power performed by the slow leg decreased in magnitude by+194 W, and peak positive power was similar across both legs for all belt speeds (Approx. 180 W).

**Figure 5.**
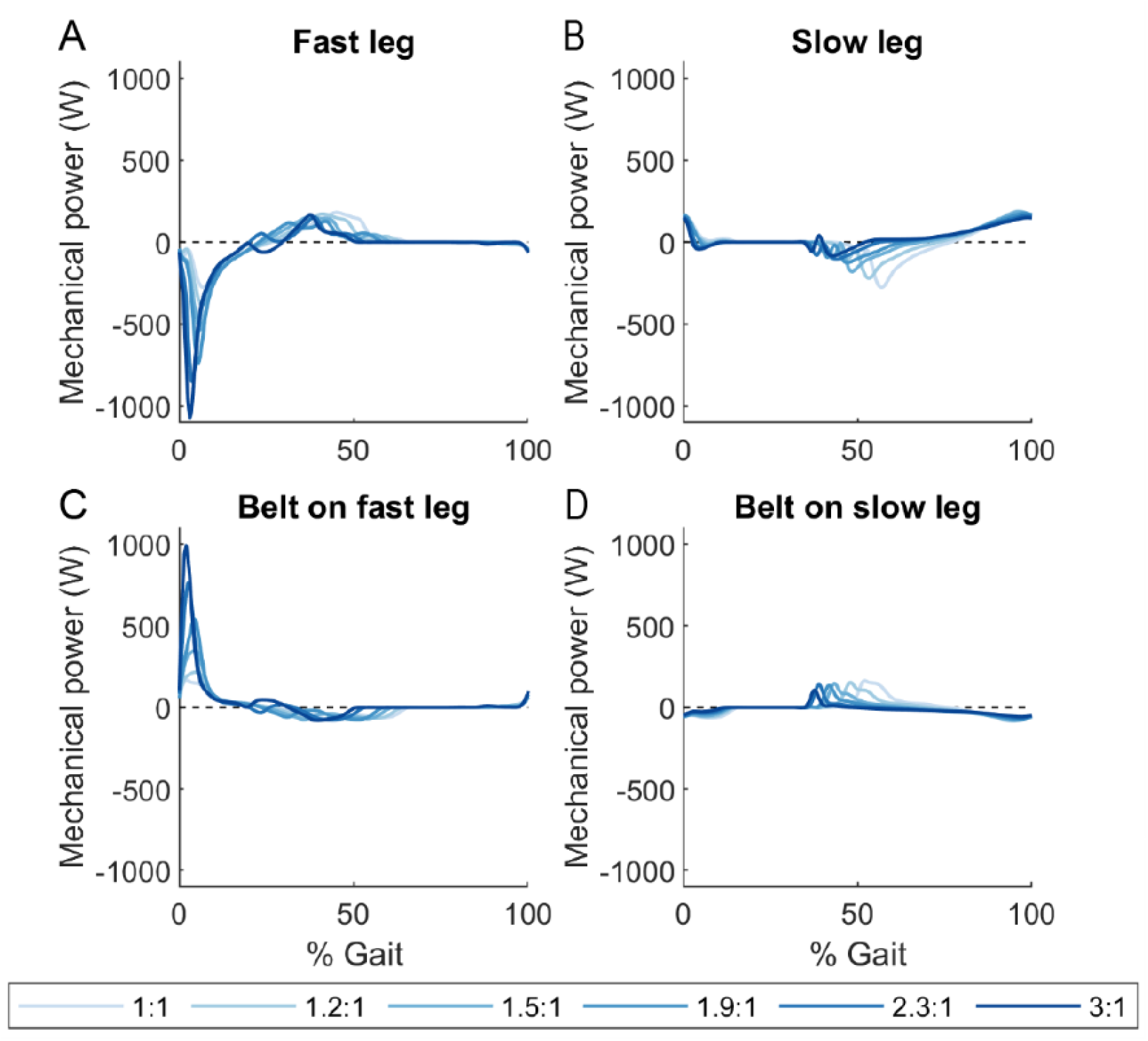
Instantaneous net mechanical power throughout the gait cycle exerted by the legs on the treadmill belts (top row) and exerted by the treadmill belts on the legs (bottom row) for belt speed asymmetry ratios ranging from 1:1 to 3:1.

The belts perform increasing amounts of net positive mechanical work on the legs as belt speed asymmetry increases (Figure 8). This change is driven by an increase in positive work performed by the fast belt on the leg of 29 W (Figure 6). The positive work performed by the slow belt on the legs decreased by 15 W, but this was mostly offset by a decrease of 12 W in negative work performed by the sum of both belts on the legs.

**Figure 6.**
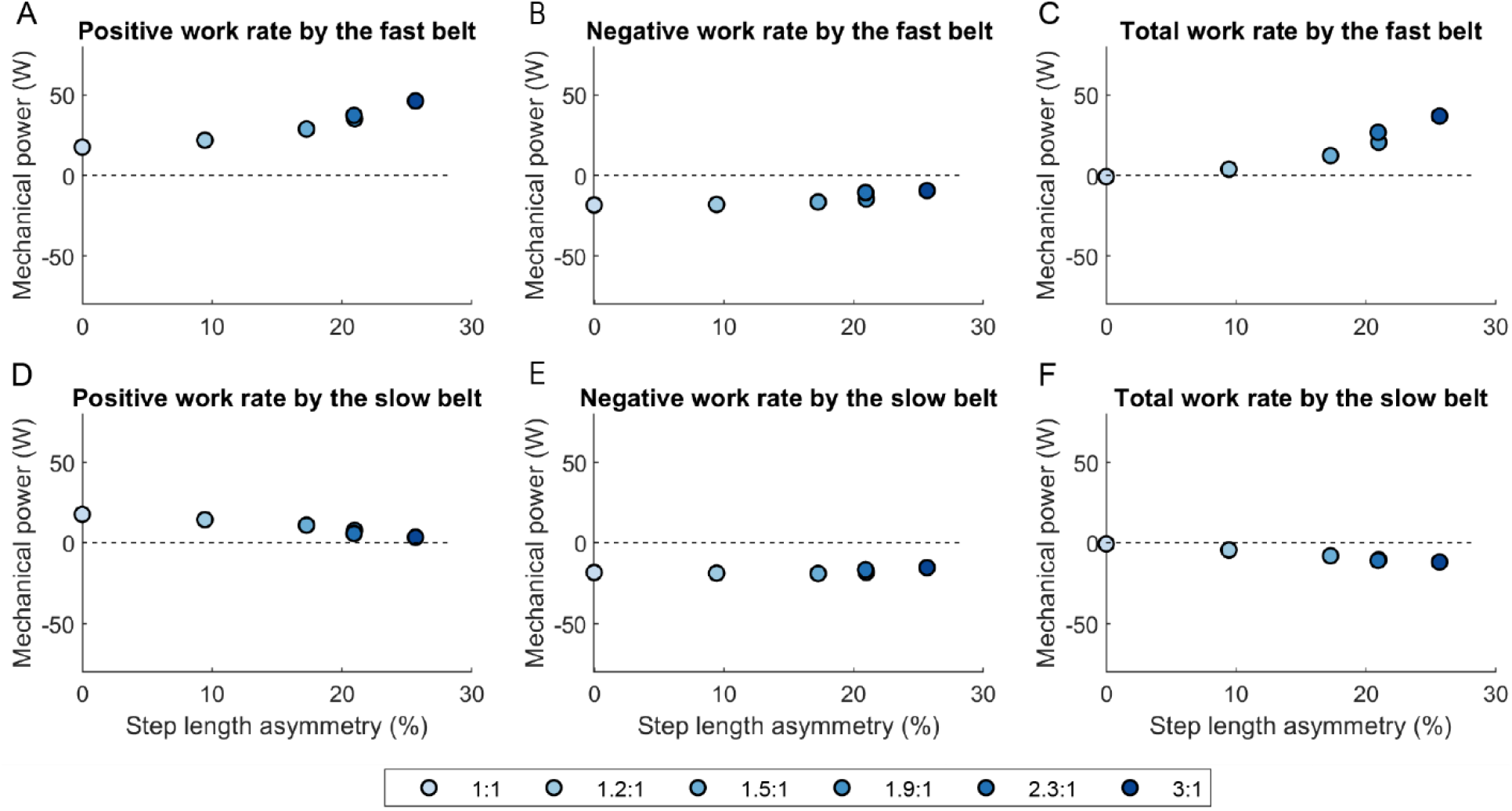
Positive, negative, and net work rate exerted by the treadmill belts on the legs vs. step length asymmetry for minimum effort gait solutions for 1:1 through 3:1 belt speed ratios, calculated per belt.

**Figure 7.**
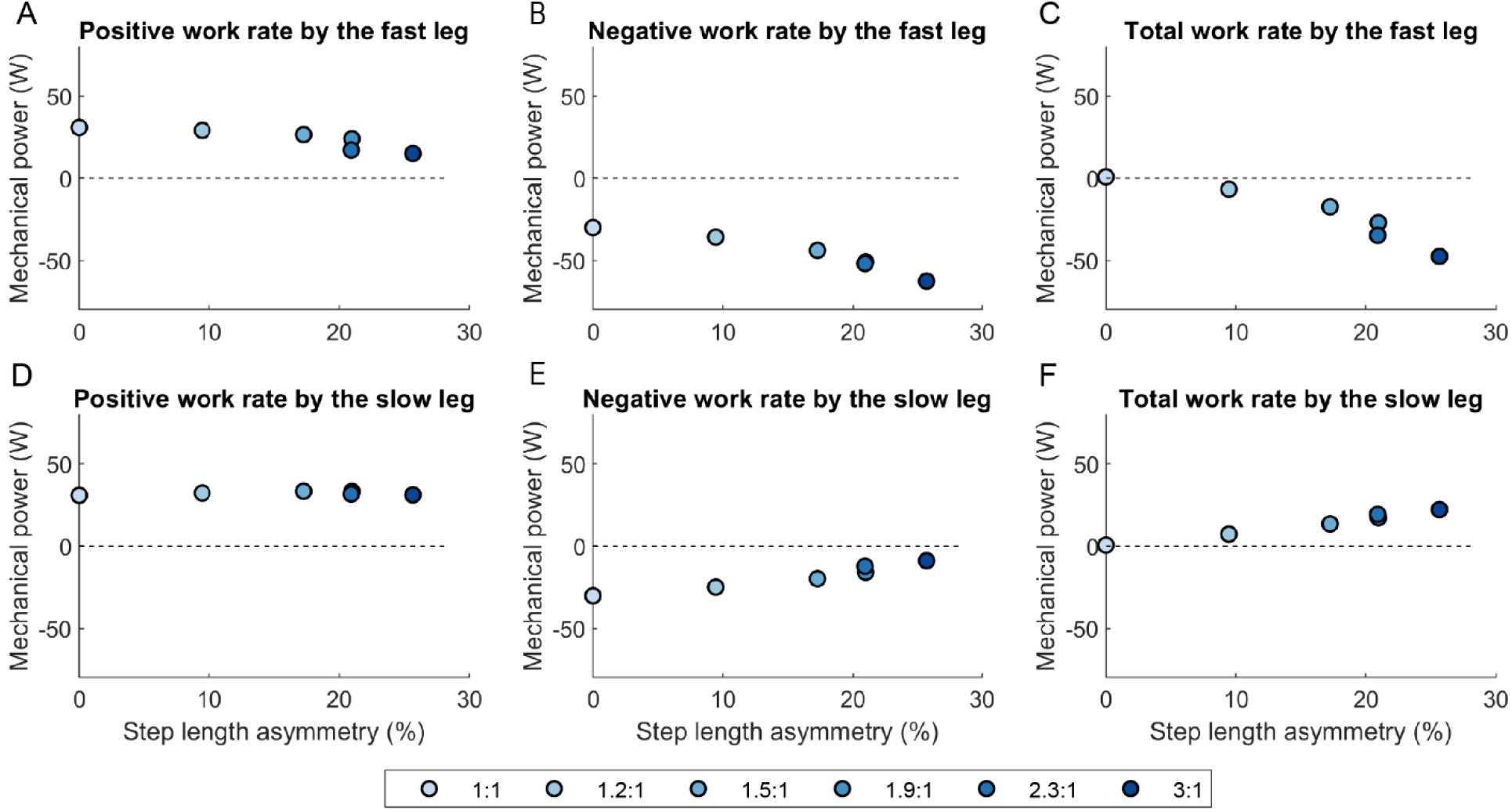
Positive, negative, and net work rate exerted by the legs vs. step length asymmetry for minimum effort gait solutions for 1:1 through 3:1 belt speed ratios, calculated per belt.

**Figure 8.**
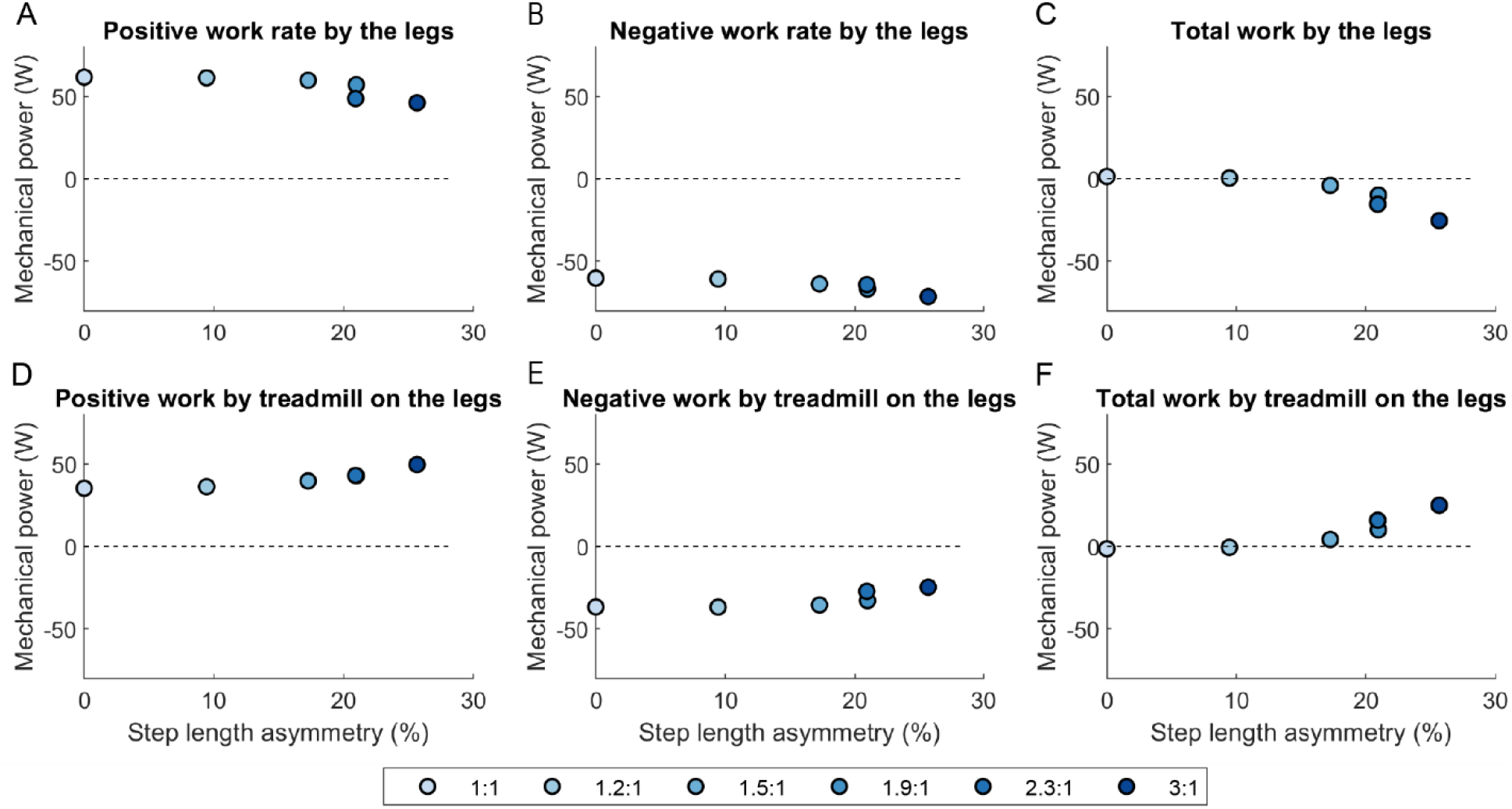
Positive, negative, and net work rate exerted by the legs on the treadmill belts (top row) and exerted by the treadmill belts on the legs (bottom row) vs step length asymmetry for minimum effort gait solutions for 1:1 through 3:1 belt speed ratios.

The legs perform decreasing amounts of net work as belt speed asymmetry increases (Figure 8). This is driven by changes in the fast leg, which had an increase in negative work of 33 W and a decrease in positive work from of 16 W for a cumulative decrease in net work of 48 W(Figure 7). These changes in the mechanical work output from the fast leg mirror and amplify the changes in fast belt mechanical work output (+37 W, Figure 6). This decrease in net work is offset by an increase of 22 W net work performed by the slow leg, which, as with the fast leg, was more exaggerated than the corresponding decrease in net work performed by the slow belt (−11 W, Figure 6). This is almost entirely accounted for by a decrease in negative work performed by the slow leg, -with little contribution via change in positive work in either direction.

These trends indicate that increasing belt speed asymmetry did not increase the mechanical energy output by the legs for propulsion on either side, but did increase the mechanical energy transferred to the legs from the belts during braking at collision, and that the model shifted the role of absorbing foot-belt collision energy toward the leg on the fast belt.

### Metabolic power

**The model exploits increasing belt speed asymmetry to reduce metabolic cost below tied-belt levels.** The model reduced the net metabolic rate by 14.3% as belt speed asymmetry increased from 1:1 to 3:1 (Figure 9a). The metabolic rate slightly increased by 0.4% as the belt speed ratio increased from 1:1 to 1.5:1, then sharply decreased as the belt speed ratio increased beyond 1.5:1 and did not appear to be approaching a lower bound at the highest asymmetry ratio simulated. This reduction in metabolic rate is linearly correlated with a decrease in positive work output by the legs, as expected (Figure 9b). The metabolic rate decreased by 1.4 W/kg for every 1.0 W/kg decrease of positive work performed by the legs (R^2^ = 0.95). Metabolic rate is inversely linearly correlated with net work performed by the treadmill on the legs (Figure 9c). The metabolic rate decreased by 0.91 W/kg for every of 1 W/kg increase of net work performed by the treadmill (R^2^ = 0.91). Correspondingly, the model reduces the positive work rate exerted by the legs in linear proportion to the increase in net work performed by the treadmill (Figure 9d). Positive work performed by the legs is reduced by 0.64 W/kg for every 1 W/kg of net work performed by the treadmill (R^2^= 0.94). This can be interpreted as the model utilizing positive net work from the treadmill to reduce positive work performed by the legs with an efficiency of 64%. This efficiency ratio has been observed to be about 33% in experiments with human participants (7).

**Figure 9.**
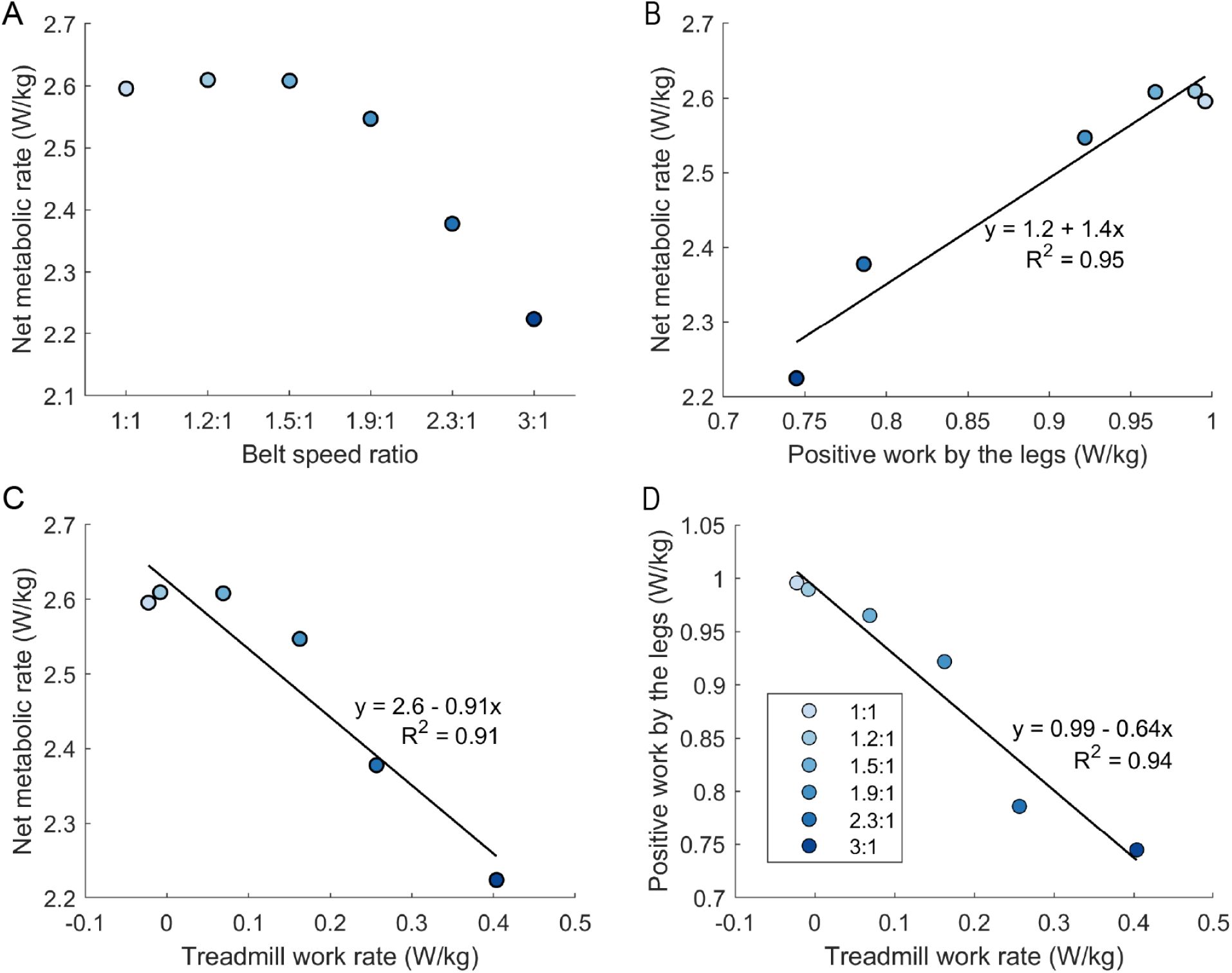
a) Net metabolic rate vs. treadmill belt speed ratio. b) Net metabolic rate vs positive work rate output by the legs. c) Net metabolic rate vs net work rate output by the treadmill. d) Positive work rate output by the legs vs. net work rate output by the treadmill. All units are normalized to the model mass.

## Discussion

This set of simulations offers a new perspective into energy-optimal biomechanics for different magnitudes of belt-speed asymmetry. They agree with physical models and experimental evidence that a treadmill with asymmetric belt speeds can exert net positive mechanical power on a walking human (7, 8). They also agree with experimental evidence that energetically optimal step length and time asymmetry increase as belt speed asymmetry increases (10). Critically, they demonstrate that the human musculoskeletal structure as modeled can exploit this mechanical power to reduce its metabolic energy consumption below the equivalent symmetrical belt speed condition. However, we have observed some key differences between the behavior of our biomechanical model and both the theoretical behavior of a simple model with pendular mechanics, as well as the experimentally observed behavior of human participants.

### Differences between optimal control simulations and human experiments

The simulated human can use the split-belt treadmill like an assistive device, as predicted by simple dynamic models. However, this result is not fully consistent with results obtained from real humans walking under similar conditions. Participants who walked with positive step length asymmetry did reduce their metabolic rate compared with negative or zero step length asymmetry when the belt speed ratio was 3:1, but not below the metabolic rate recorded during the symmetrical belt speed condition (7, 8). Likewise, local energetic minima were found at positive step length asymmetry values for three belt speed differences, but the metabolic energetic cost at each minimum increased with greater belt speed asymmetry in (Stenum & Choi, 2020). Experiments across a wide range of belt speed combinations found that the metabolic cost of walking on asymmetric belt speeds was 17.5% larger on average than the cost of walking on tied belts (11).

Additionally, the optimal step length asymmetry is greater for the model than it is in human experiments. Consistent with the simulated model, energetically optimal step length asymmetry was observed to increase with increasing belt speed asymmetry in (10). However, the model optimum for a 3:1 belt ratio (+0.256) was notably greater than the largest step length asymmetry optimum observed experimentally with 4:1 belt speed ratio (+0.208, when measured as the distance between ankle markers at heel strike. In (7), optimal step length asymmetry was estimated to be between +0.06 and +0.38, though only two participants walked at target asymmetries above +0.15 and their metabolic rate at these asymmetries was greater than the predicted minimum for both.

One explanation as to why our simulations walk with greater step asymmetry and reduce the metabolic power below tied-belt walking is that the model can exploit the mechanics of the split belt treadmill much more efficiently than humans have demonstrated. It is possible that simplifications in the biomechanics can account for some of this difference; mechanical energy transmitted from the belts is constrained to the sagittal plane, and no energy is dissipated within the joints themselves when the foot collides with the ground. Additionally, our choice of contact model may affect braking and propulsion power at heel-strike and toe-off.

The model’s superior efficiency can also be explained, however, by the lack of any control objectives aside from muscle effort minimization. One way in which this is evidenced is by the magnitude of instantaneous mechanical power transmitted to the legs from the treadmill upon heel-strike on the fast belt. In the current problem formulation, the model experiences no penalty for experiencing extreme shock loads and will adjust its behavior to ensure that they occur if they can reduce the required amount of muscle activity to perform locomotion by even a marginal amount. Similarly, a human walking with the model-optimum level of asymmetry may perceive it to be unstable, which the model does not account for. Humans do not necessarily choose energetically optimal gait patterns when doing so would decrease stability (e.g., walking downhill) (30). Additionally, it is possible that the human nervous system contains biases toward step length symmetry as a consequence of the underlying neural circuitry or higher-level motor control objectives that compete against energy optimization.

While many studies have observed how humans adapt to walking on split belt treadmills, none have specifically investigated long term adaptation of participants who are additionally constrained to maintain a substantial positive step length asymmetry. Training can have a significant effect on the outcome of interacting with an assistive device; for example, the type and duration of training had a significant effect on the metabolic cost of transport of people walking with exoskeleton assistance (31). Thus, it is also possible that humans did not walk with such extreme step length asymmetry because more training was needed to develop efficient motor control strategies that take full advantage of the mechanical benefits (e.g. excessive co-contraction due to task unfamiliarity).

### Differences between optimal control simulations and simple dynamic model

Our simulations reproduce patterns predicted by simpler models with pendular mechanics, but do contain some distinctions. Positive SLA necessarily results in net positive work performed by the treadmill on an inverted pendulum walker which exerts active forces on the ground via piston-like compressible legs and a single contact point at each foot. The musculoskeletal model we used more closely resembles the dynamics of the human lower limb, and while pendular dynamics apply during early stance, the straight compressible leg approximation does not hold for mechanical energetics calculations during late stance.

The main reason for this difference is that a typical human leg can both exert a propulsion force with ankle plantarflexion regardless of limb angle and lift itself into the swing phase without pushing off against the ground by flexing the knee and hip. Initiating the swing phase without substantial ground push-off is not an efficient gait strategy during normal walking, but it becomes advantageous when there is a strong energetic penalty associated with pushing off against the ground on one side. To balance the anterior-posterior impulse and keep the model from drifting backward on the treadmill, the simulation also reduces the collision angle of the leading leg by placing the foot closer to the center of mass, as expected if governed by pendular mechanics during early stance.

This strategy effectively removes the role of the trailing foot position from any gait adjustment to reduce propulsion force on the fast belt or increase it on the slow belt. It is likely that any bipedal gait model would struggle to reduce its muscle activity relative to the tied belt condition if this were not the case, because the belt speed difference makes a shallow trailing limb angle on the fast belt difficult to achieve at high belt speed ratios without extreme differences in step time that may require disadvantageous mechanics during swing.

Another major difference is that human walking gait has a double-support phase, causing the trailing limb to typically output its peak propulsion force after the moment that step length is measured. For a model with a double-support phase, the mechanical work calculations require accounting for trailing limb hGRF after heel-strike of the opposing leg. However, because our simulations produced a positive SLA that expressed mainly with leading foot placement differences, which was the spatial measure that drove most of the mechanical work changes between the human and the treadmill, positive SLA was still correlated with positive net mechanical work performed by the treadmill. Step length, however, does not appear to directly drive the flow of mechanical energy from the treadmill to the legs, but rather a combination of leading foot placement at heel-strike and muscle control strategy during late stance.

These differences can be accounted for by simplifying assumptions in the simple pendular model. However, we observed that the mismatch between human behavior and the theoretically optimal behavior predicted by this model is persistent with a more complex model, suggesting that the difference is because humans do not optimize their behavior to minimize effort at all costs.

### Modeling considerations and limitations

Aside from modeling assumptions acknowledged above, these simulations have limitations that may be the subject of future work. First, our optimization framework is not able to generate a cost landscape, making the energetic consequences of step length symmetry changes for a constant belt speed ratio impossible to assess. Our simulations produce less asymmetry for lower belt speed ratios, implying that energetic gains from increasing step length asymmetry are bounded. Enforcing specific step length or step time asymmetry values, as implemented for overground walking in (32), would allow optimal control simulations to generate cost landscapes for comparison with experimental studies.

Second, our simulations do not include a systematic distribution of limb length or mass to represent a participant population. It is possible that the optimal gait strategy might vary for humans of different sizes. This limitation could be addressed in future by simulating an array of virtual participants with varying physical parameters, as done in (20). To mitigate our own concerns about drawing strong conclusions from the output of a single model, we additionally performed the same set of simulations for a model with additional degrees of freedom (namely: MTP joints, additional muscles in the foot, and implicit tendon dynamics) adapted from the model used in (32) to verify that the trends from our results are not affected by our most basic model simplifications. This model has a 21% larger mass and approx. 3% longer limbs with different values for the maximum isometric force for nearly every muscle. While not a systematic set of model variations, we did find that all of the trends in our reported results were exhibited by both models. The full list of parameters for this model (Table S1) and set of results (Figures S2-S9) are included as supplemental information.

Finally, our choice of muscle specific tension affects the magnitude (but not the trend) of the relationship between mechanical work and metabolic rate. The specific tension of human skeletal muscle does not have a single agreed-upon value, requiring computational models to assign an estimate which generates realistic joint moments and metabolic energy costs (28, 33). Our value of 60 N/mm^2^ has been used in other muscle-driven simulation studies which meet these criteria (20, 33), but the amount of mechanical work the model is able to convert into metabolic energy savings may vary in real human gait, even with optimal mechanics. This does not influence the relationship between mechanical work performed by the treadmill and the human model, however.

### Implications for human experiments

These simulations demonstrate gait behavior when muscle effort is minimized, necessarily driving the behavior toward exploiting the passive dynamics of the system and constraints. Stable gait using only passive mechanics cannot be achieved on a level split-belt treadmill with typical human anatomy, but optimal control simulations can find gait patterns that require minimal added torques from muscle activity. These patterns may be sufficiently different from neuromuscular control patterns for gait to require training akin to learning a new motor skill before participants will be able to use belt-speed asymmetry to reduce the metabolic energy cost of walking below tied-belt levels. On the other hand, energy-optimal gait behavior with asymmetric belt speeds may be harmful to the body at asymmetry magnitudes large enough to make a significant difference to the energy cost of walking, which may bound the degree to which humans will normally exploit belt speed asymmetry even if their gait strategy is driven primarily by energy optimization.

Optimal control behavior provides a reference point for experiments to test these ideas. Assessment of adaptation strategy in split-belt studies may be improved by observing critical measures in optimal control solutions such as fast-belt braking force magnitude rather than focusing exclusively on step length or time symmetry. Evaluation of joint loading with respect to asymmetry measures, foot placement, or ground reaction force asymmetry in human experiments is critical to understanding the bounds on observed behavior aside from symmetry error or energy cost objectives. Additionally, results from future experiments may compare our optimal control solutions with expected outcomes from simple pendular models to determine if the added complexity of a full musculoskeletal model better explains experimental trends.

Finally, it is possible that if split belt treadmill adaptation is primarily driven by exploiting the system dynamics to reduce energy expenditure, the transference of these adapted gait strategies to situations where the dynamics cannot be exploited in the same way may be limited (i.e. overground walking). This could have an impact on the efficacy of treadmill-based motor adaptation protocols for rehabilitation. Reproducing treadmill-based motor learning experiments overground is important to determine whether adaptations can be reproduced without the unique constraints of a treadmill.

## Conclusion

The simulated results agree with the experimental finding that walking with positive SLA on asymmetrical belt speeds results in positive net work performed on the human by the treadmill. Simulations converged to much higher magnitudes of SLA than humans chose even after experiencing high SLA gait patterns, and they also reduced the metabolic rate from the tied-belt condition. This simulation framework presents a scenario for what human gait might look like if maintaining steady-state walking with the least possible amount of muscle effort were the only goal of locomotion. We interpret the overall reduction of metabolic rate and the dramatic increase in SLA compared to experimental observations to mean that the model is able to take advantage of the asymmetry of the split-belt treadmill to reduce the energy cost of walking more efficiently than humans so far have managed. This may be due to factors such as the model’s lack of concern for stability or joint loading, the potentially insufficient adaptation time to achieve mastery of optimal biomechanics in prior studies, and model simplifications.

## Availability of data and materials

Supplemental tables and figures are available at: https://figshare.com/s/f22e98068d83a2fd031d

All models and code are available at https://simtk.org/projects/split-belt-gait

